# TNF controls a speed-accuracy tradeoff in the apoptotic decision to restrict viral spread

**DOI:** 10.1101/2020.02.20.958942

**Authors:** Jennifer Oyler-Yaniv, Alon Oyler-Yaniv, Evan Maltz, Roy Wollman

## Abstract

Early commitment to apoptosis is an important antiviral strategy. However, fast decisions that are based on limited evidence can be erroneous and cause unnecessary cell death and tissue damage. How cells optimize their decision making strategy to account for both speed and accuracy is unclear. Here we show that exposure to TNF, which is secreted by macrophages during viral infection, causes cells to change their decision strategy from “slow and accurate” to “fast and error-prone”. Mathematical modeling combined with experiments in cell culture and mouse corneas show that the regulation of the apoptotic decision strategy is critical to prevent HSV-1 spread. These findings demonstrate that immune regulation of cellular cognitive processes dynamically changes a tissues’ tolerance for self-damage, which is required to protect against viral spread.

## Main

Rapid detection of viral infection is needed to minimize spread and associated damage. At the same time, evolutionary pressures constantly enhance the arsenal of strategies that viruses use to evade and manipulate host detection ^1,2^. The adversarial co-evolution between host detection and viral avoidance results in a challenging and time-dependent classification problem for host cells that likely operate close to the detection limits imposed by the inherent stochasticity of biochemical reactions. Rapid decision-making at the noise limit is likely to result in false-positive classification i.e., a decision that a virus is there when it is not, followed by unnecessary cell death and bystander tissue damage. How cells navigate the tradeoff between decision speed and accuracy to achieve an optimal classification strategy has not been explored.

The initial detection of viral invasion in a tissue is accomplished by macrophages and other innate immune sentinels ^3^. These specialized cells secrete cytokines and other inflammatory mediators to alert nearby cells of invading pathogens. Alerted cells then execute diverse strategies to prevent viral spread including restriction of viral entry, direct inhibition of viral gene expression, prevention of viral genome replication, prevention of viral egress, and apoptosis of infected cells ^4,5^. These defence mechanisms are activated in an inducible manner as they have the potential to interfere with normal host cell physiology ^6–9^. We asked how cells balance the competing goals of antiviral activity and maintenance of normal physiology in response to macrophage-derived inflammatory cytokines. We focused on Tumor Necrosis Factor ɑ (TNF), because it is an important proinflammatory cytokine, yet can also cause significant damage to healthy host tissues due to its role regulating the apoptotic pathway ^10^. This relationship is exemplified by the observation that TNF-blocking drugs have shown remarkable success for treatment of autoimmune disorders, yet also increase susceptibility to certain pathogens ^11^.

### TNF secretion by Macrophages limits viral spread

We first quantified the effect of macrophage density, inflammatory status, and TNF secretion on viral spread throughout a population of cells. We co-cultured fibroblasts with either naive or activated mouse bone marrow-derived macrophages (BMDM) at different densities and infected them with a low multiplicity of infection (MOI 1) of Herpes Simplex Virus-1 (HSV-1). We used HSV-1 recombinant viruses expressing capsid protein VP26-fluorescent protein fusions to monitor viral infection using time-lapse microscopy ^12^. As expected, activated, but not naive, BMDM restricted viral spread (Fig 1A-B, movies S1-S4). Activated macrophages produced high levels of TNF (Extended data 1A), suggesting that this cytokine is a potential key regulator. Indeed, BMDMs derived from mice that lack TNF (TNF KO), and BMDMs supplemented with TNF neutralizing antibodies did not restrict viral spread (Fig 1A-B, Extended data 1B). These results show that the restriction of viral spread by inflammatory macrophages depends on the production of TNF. Interestingly, the protective effect of TNF against viral spread was not mediated by reduced cell infectivity or by enhanced sensing of HSV-1 or synthetic viral ligands (Fig 1C-E, Extended data 1C-E).

**Figure 1:**
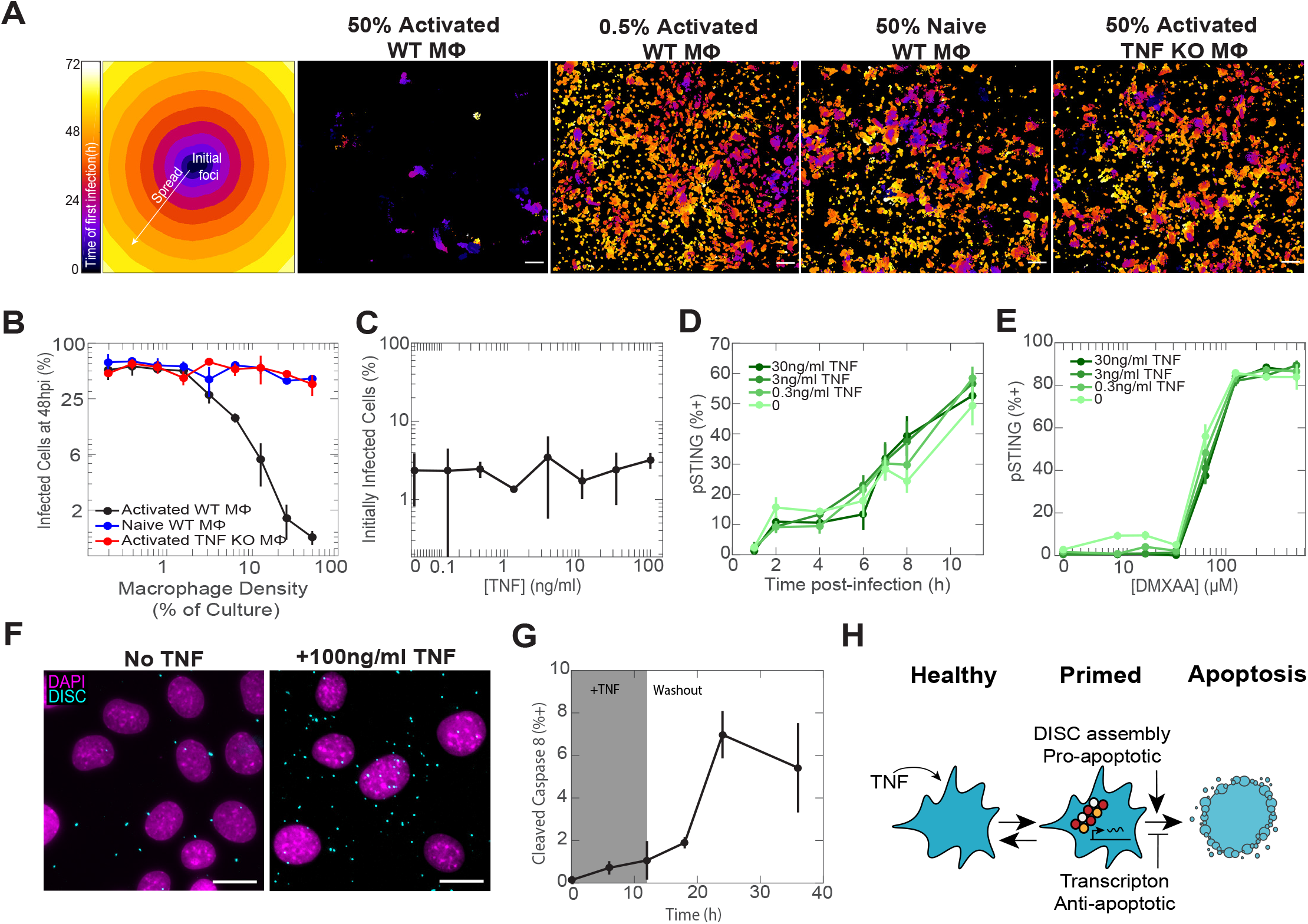
Activated macrophages restrict viral spread by production of TNF. **A**, Representative temporal color-coded maps of viral first infection time for selected BMDM-fibroblast co-culture conditions (scale bar, 100μm). **B**, Quantification of viral spread across all BMDM-fibroblast co-culture conditions. **C**, Quantification of cells initially infected only from virus added to the media after being pulsed overnight with TNF. **D**, Quantification of cells that stained positive for active, phosphorylated STING (pSTING) after being pulsed overnight with TNF and infected with MOI 10 of HSV-1. **E**, Quantification of cells that stained positive for pSTING after being pulsed overnight with TNF and exposed to dose titrations of the pSTING agonist DMXAA. **F**, Representative images of cells assayed for DISC assembly via Traf2-Caspase 8 association (scale bar, 30μm). **G**, Quantification of cells that stained positive for cleaved Caspase 8 during and after TNF treatment. The shaded region indicates the time period before TNF was washed out. **H**, Cartoon depicting the temporary primed-to-death cell state induced by TNF and characterized by DISC assembly and initiation of a pro-survival gene expression response. Data in **B-E** and **G** are mean±s.d.

We next investigated whether viral defense is related to TNF augmentation of the cellular propensity to undergo apoptosis (Extended data 2, 4A). Exposure to a saturating dose of TNF (100ng/ml) for 6 hours showed only a minimal increase in cell death (Extended data 3A). However, since TNF simultaneously activates opposing pro-apoptotic and pro-survival signals ^13–17^, we reasoned that activation of apoptotic signaling can occur without a corresponding increase in cell death. Consistent with that, co-treatment of TNF and Actinomycin D, which blocks transcription, resulted in the rapid death of nearly the entire cell population ^10^ (Extended data 3A). We then quantified the effect of TNF on the formation of Death Inducing Signaling Complex (DISC) ^18^, which controls downstream activation of initiator caspases ^19–21^ (Fig 1F, Extended data 3B-C). We saw a dramatic increase in the population frequency and per-cell abundance of DISC assembly in cells treated with TNF compared to untreated cells. Although TNF induced a majority of cells to assemble DISC, only a small fraction activated caspase cleavage (Fig 1F-G, Extended data 3B-C).

We further identified the apoptosis inhibitors *ciap1* and *ciap2* as key components of the pro-survival transcriptional response (Extended data 3D). Subsequently, pharmacologic inhibition of cIAP1 and cIAP2 with LCL-161 ^22^ induced widespread apoptosis in cells primed with TNF, but not in untreated cells (Extended data 3E). Finally, to monitor the dynamic nature of the increase in apoptotic propensity we measured cleaved caspase-8 after TNF was washed and found a sustained increase in cell death that persisted for at least 36 hours post TNF treatment (Fig 1G, Extended data 3F). Collectively, these data illustrate that TNF transitions cells into a reversible, ligand-independent “primed-to-death” cell state in which DISC formation constitutes a trigger that is counteracted by pro-survival proteins (Fig 1H).

### Speed vs accuracy tradeoff

Rapid apoptosis following viral detection is highly advantageous ^5,23^ and therefore we hypothesized that the TNF-induced “primed for death” cellular state allows cells to accelerate their commitment to death. To measure the speed of cell decision, we treated fibroblasts with dose titrations of TNF, infected them with MOI 10 of HSV-1, and imaged viral infection and cell death for 48 hours. MOI 10 causes a synchronous and uniform infection minimizing the impact of viral spread which simplifies interpretation. We tracked individual cells and identified the timing of infection and death (Fig 2A). We observed that TNF treatment shortened the time interval between infection and death. TNF did not shorten the time from the increase in caspase-8 activity to death, which was much shorter than the time from infection to caspase-8 activation ^24^ (Extended data 4A-C). We will refer to the time interval between infection and death as the ‘apoptotic decision latency’. To further establish that the decision latency is a rate-limiting step we fitted the measured single cell decision latencies to an exponential distribution and found that the average latency matched the rate of population decline (Fig 2B-C, Extended data 4D, F). From these population measurements, we concluded that TNF dose-dependently shortens the apoptotic decision latency (Fig 2D).

**Figure 2:**
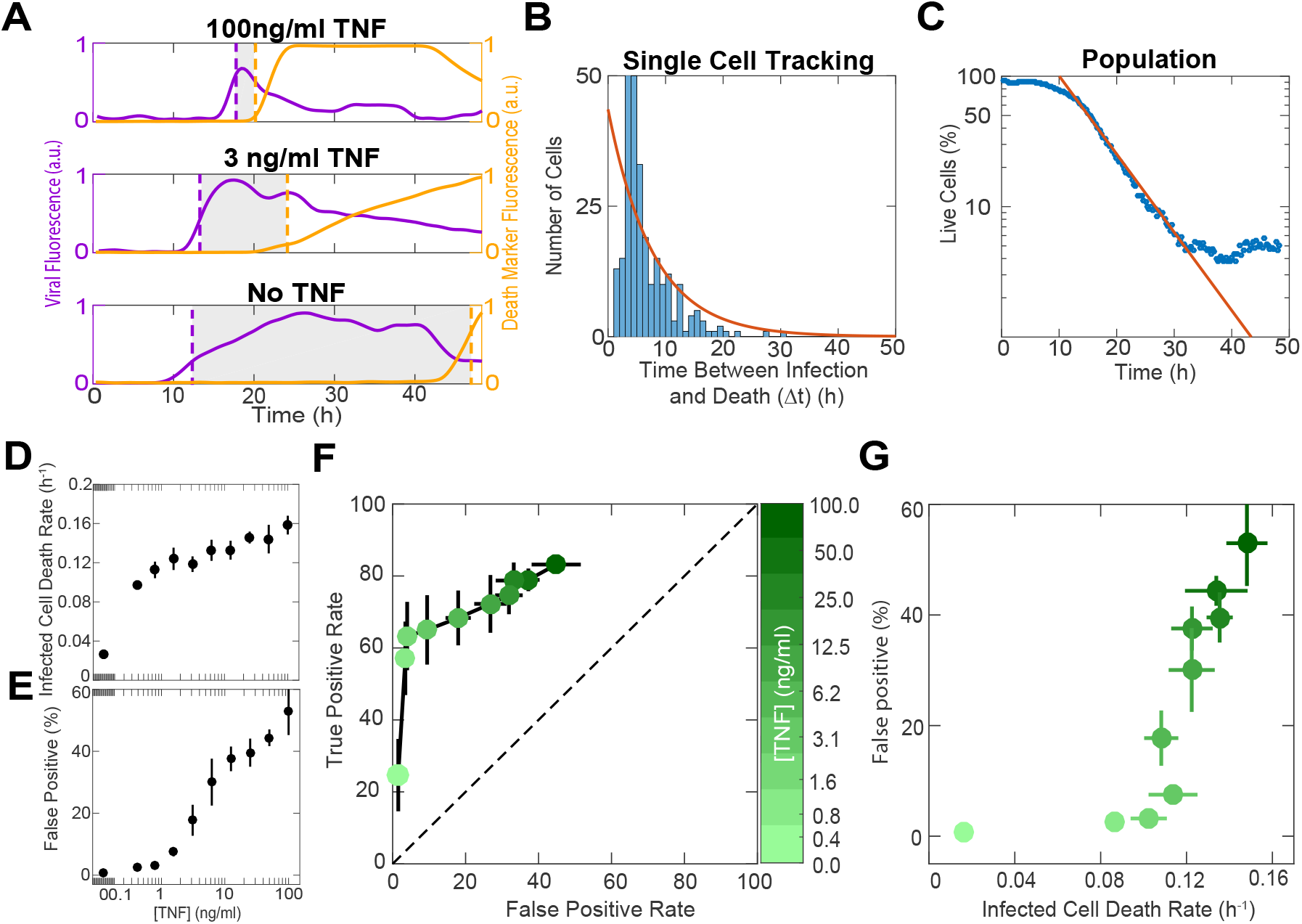
TNF regulates the tradeoff between apoptotic decision speed and accuracy. **A**, Representative single cell traces of cells treated with different doses of TNF and infected with MOI 10 of HSV-1. Dotted purple and yellow lines indicate the timing of initial infection and death, respectively. Shaded regions represent the time interval between infection and death (apoptotic decision time, Δt). **B**, Distribution of apoptotic decision times obtained by single cell tracking of cells treated with a single dose of TNF and infected with MOI 10 of HSV-1. Orange line shows an exponential fit of the data. **C**, The same sample that generated **B**, plotted as the percentage of live cells over time. Orange line shows the same exponential fit plotted in **B**. **D**, Death rates obtained from exponential fits of cells treated with dose titrations of TNF and infected with MOI 10 of HSV-1. **E**, Quantification of the percentage of uninfected cells killed by TNF (false positives) at 24 hours post treatment. **F**, ROC curve where false positive rate represents cells killed by TNF at 24 hours post treatment and true positive rate represents infected cells that die during the first viral life cycle (10 hours after initial infection). **G**, Speed versus accuracy tradeoff shown by plotting the percentage of false positive cells (from **E**), versus the death rate for infected cells (from **D**). **D-G** are mean±s.d. **F,G** colorbar shows TNF concentration.

Our data show that TNF dramatically shortens the decision latency following viral infection. We asked if there are downsides to this short-latency state that constrain the speed of cellular decision making. A tradeoff between decision speed and accuracy is well established in other cognitive systems ^25,26^ we therefore analyzed whether the increasing speed in commitment to apoptosis comes at a cost of accuracy. Two possible decision errors are false negative - infected cells that do not die - or false positive - cells that die without being infected. In our MOI 10 experiments, there were almost no infected cells that escaped death (false negatives), by 48 hours post-infection (hpi) (Extended data 4D). On the other hand, the rate of false positive errors increased in a dose-dependent manner with increasing concentration of TNF (Fig 2E, Extended data 4E). Using a receiver operator curve (ROC), a standard assessment of a classifier quality, we saw a tradeoff between accurate decisions (true positives) and erroneous ones (false positives) (Fig 2F). Contrasting the increase in error rates with the increase in decision speed reveals that the cellular decision to commit apoptosis is subject to a tradeoff between speed and accuracy (Fig 2G). Our data therefore demonstrate that the cell fate decisions are not only subject to the speed-accuracy tradeoff but that this tradeoff is tunable and regulated.

### Regulation of speed-accuracy tradeoff prevents viral spread

We next investigated how the dynamics of viral spread are affected by TNF modulation of the apoptotic decision strategy. We confirmed that TNF restriction of viral spread depends on the apoptotic pathway (Extended data 5A-B) supporting that viral spread depends on the interplay between infection and death. To better understand this connection we used a stochastic and spatially explicit mathematical model of viral spread. In our model, each cell can be in one of four states: healthy, infected, dead following infection, or dead but uninfected (Fig 3A). The cells are organized on a uniform triangular grid and the simulation is initialized by infecting the central cell on the grid. These conditions are analogous to a low MOI infection where a small fraction of cells form an initial seed from which the virus spreads radially in a juxtacrine manner. The viral load of each infected cell grows linearly and deterministically while the cell is alive (Extended data 5C). Healthy cells can be infected by their infected neighbors and the probability of infection is determined by the total viral load of a target cell’s live neighbors (Extended data 5D). The death rates of individual cells depend on their infection status, their exposure to TNF, and are directly interpolated from measured values (Extended data 5E-F). All model parameters are based on individual cell decisions and were independently calibrated in separate experiments (Extended data 5C-F).

**Figure 3:**
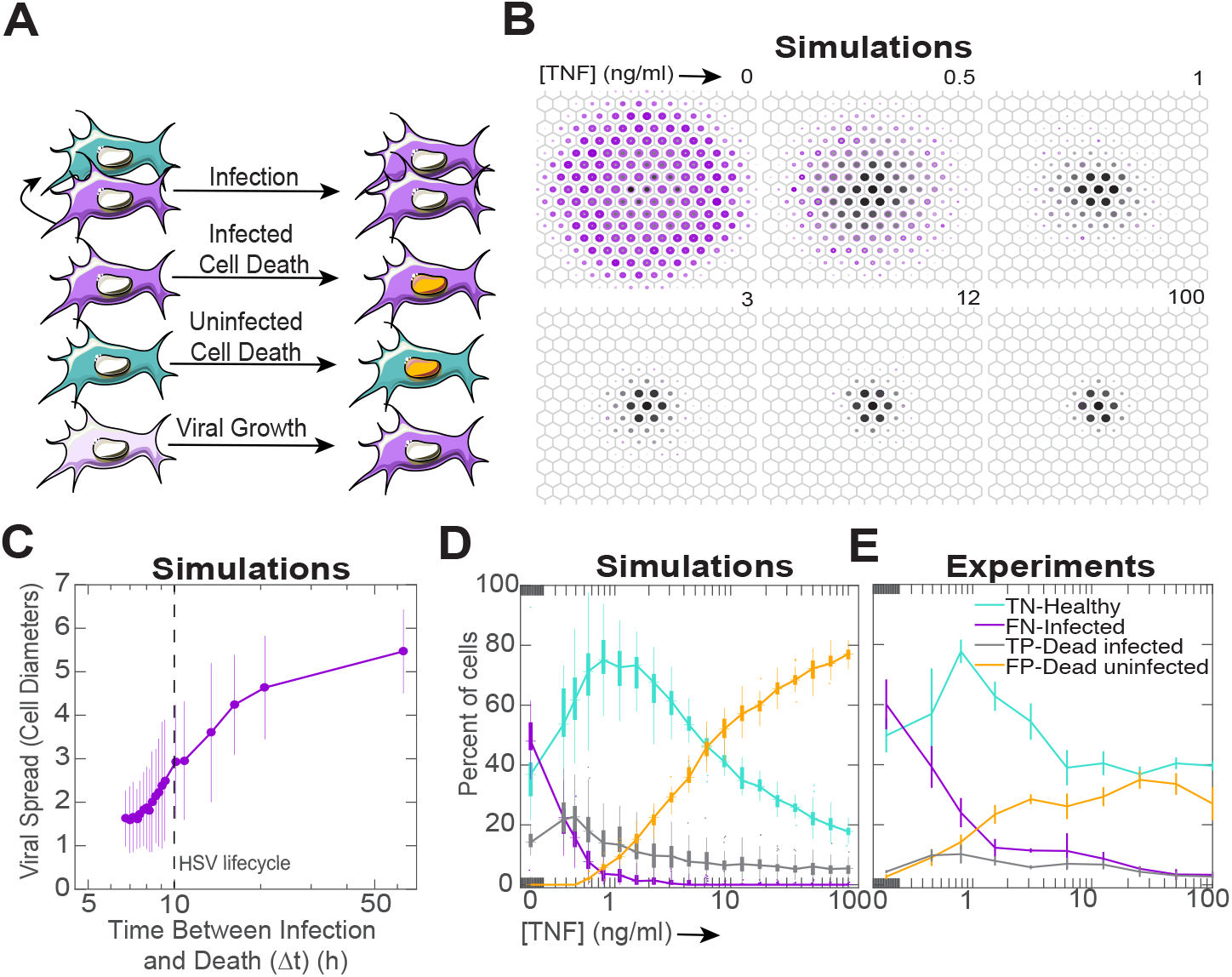
Modulation of the tradeoff between cellular apoptotic decision speed and accuracy controls viral spread throughout a population. **A**, Diagram of the reactions in our spatial stochastic model. All model parameters were measured in experiments shown in extended data 5b-e. **B**, Ensemble simulation results for different TNF concentrations. Purple area and intensity correspond to the probability of the cell being infected. Black area and intensity correspond to the probability of the cell being dead following infection. **C**, Quantification of viral spread as a function of TNF concentration, mean±s.d. **D**, Boxplot summary of simulation results for a dose titration TNF conditions showing the percentages of healthy, infected, dead following infection, and dead but uninfected cells. **E**, Quantification of the percentages of healthy, infected, dead following infection, and dead but uninfected cells treated with dose titrations of TNF and infected with MOI 1 of HSV-1. For simulations (**B-D**), each condition was simulated 50 times.

Our model demonstrates that the TNF-mediated manipulation of apoptotic rates on a single-cell level is sufficient to explain the emergent arrest of viral spread (Fig 1A-B, Fig 3B, D-E). Our results show a sharp system-wide transition from a virus-permeable to a virus-resistant state which occurs when the single-cell apoptotic decision time dips below the viral life cycle ^27^ (Fig 3C). The model also captures a global manifestation of the speed-accuracy tradeoff: it predicts a sweet spot of TNF concentrations that optimizes the balance of costs and benefits (Fig 3D). This prediction was validated by infecting fibroblasts with a low MOI of HSV-1 in the presence of different doses of TNF and observing a similar sweet spot (Fig 3E).

### Prevention of viral spread in the cornea

We next sought to establish that TNF regulation of cellular decision strategies is sufficient to alter viral spread in a whole live organ. To this end, we infected mouse corneas, dissected from R26-H2B-mCherry mice, with fluorescent HSV-1 (Fig 4A). The cornea is an ideal model for our experiments: it is transparent, avascular, and is very robust in organ culture conditions. Biologically, it is a natural target of HSV-1 ^28^, and key to these experiments, it has minimal innate immune activity once dissected from its source of recruited immune cells ^29^. In our experiment we took advantage of this fact and compared corneal response with and without the addition of TNF to the organ culture media, mimicking an impact of recruited inflammatory macrophages ^30^. Using a custom light sheet microscope (Extended data 6) we imaged viral spread and cell death continuously over 48 hours. Time-lapse imaging revealed that HSV-1 spread radially over time and created a pronounced viral nodule containing an inner core of dead cells (movie S5, Fig 4C). Viral spread was dramatically reduced in corneas treated with exogenous TNF compared to untreated controls (movie S6, Fig 4B-D). In addition, by 48hpi, almost all virus-infected cells were dead in the TNF-treated conditions compared to only about 50% in untreated corneas, demonstrating that viral spread outpaced cell death (Figure 4C, E). Using dissociated freshly-isolated corneal epithelia, we validated that the increase in cell death among infected cells stems from a TNF induced increase in decision speed (Fig 4H), in agreement to our observations in fibroblasts (Fig 2G). Finally, and similar to our *in vitro* observation, TNF without infection increased the amount of bystander cell death in the cornea, fully recapitulating the apoptotic decision speed vs accuracy tradeoff (Fig 4F-G). Taken together, these data demonstrate that TNF modulates the apoptotic decision strategy of corneal epithelial cells to restrict viral spread *in vivo*.

**Figure 4:**
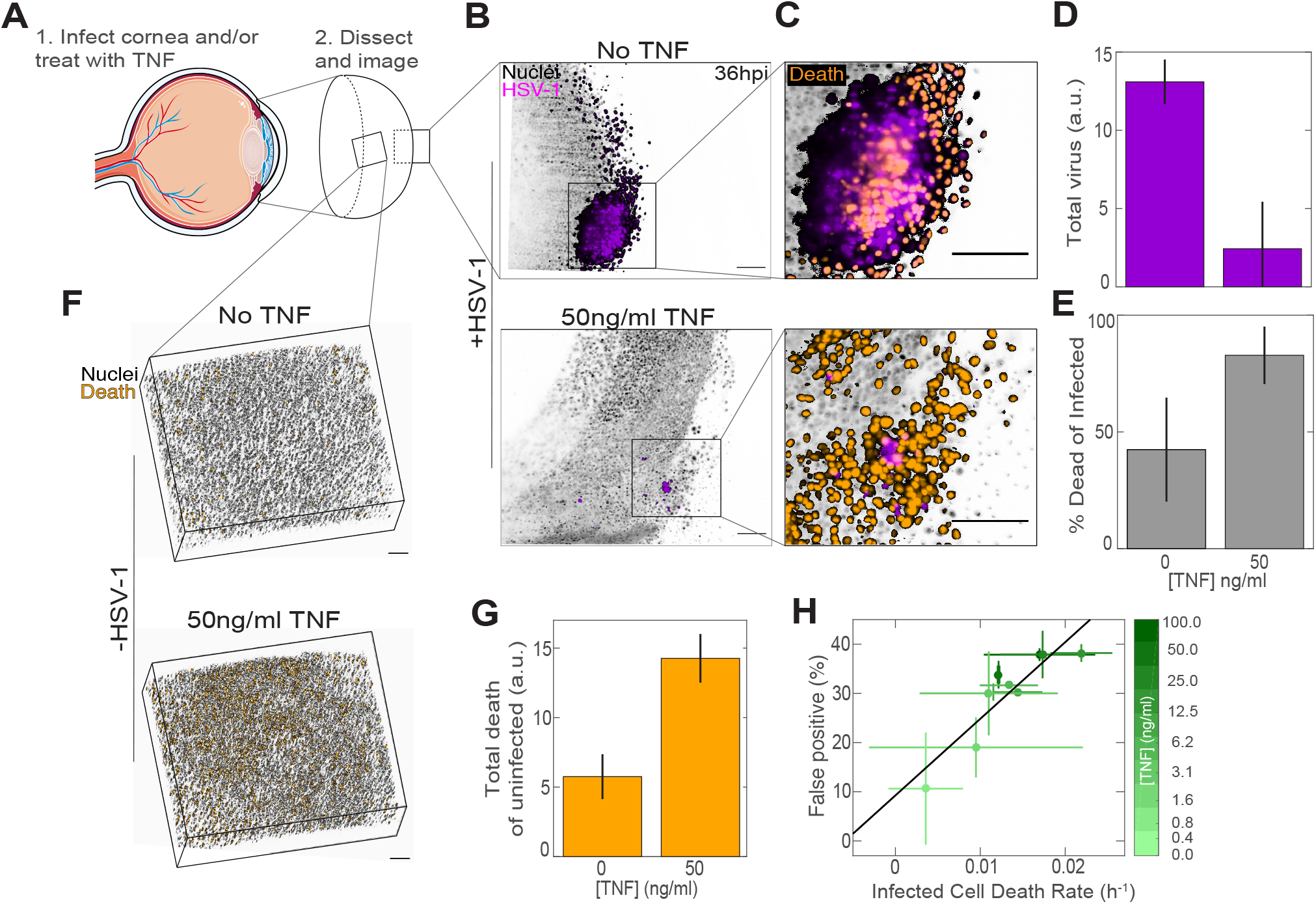
TNF alters the cellular apoptotic decision-making strategy in the corneal epithelium, which restricts viral spread. **A**, Cartoon diagram with cornea emphasized to show different perspectives of images. **B**, Representative transverse image of R26-H2B-mCherry mouse corneas infected with HSV-1 and treated with either 0 or 50ng/ml TNF. **C**, Enlarged regions within images shown in **B** with cell death channel included. **D**, Quantification of the total sum of virus pixels for C57Bl/6 corneas treated with 0 or 50ng/ml TNF, infected with HSV-1, and imaged at 48hpi. **E**, Percentage of total virus pixels also positive for Sytox Green death marker. **F**, Representative epithelial projections of C57Bl/6 corneas treated with 0 or 50ng/ml TNF and imaged 24 hours after treatment. **G**, Quantification of the total Sytox Green intensity for uninfected corneas. Data in **D-E, G-H** are mean±s.d. All scale bars, 100μm.

## Discussion

The data presented here demonstrate that information provided to cells through macrophage secreted cytokines can alter their classification strategy by dynamically shifting their error tolerance in favor of faster decisions. Importantly, these changes to individual cells’ decision processes are required to prevent viral spread in the population.

We only observed a TNF induced change in cellular decision strategy during viral infection and not in response to other stresses such as bacterial lipopolysaccharide or DNA damage (Extended data 7). One interpretation of this context specificity stems from the fact that viruses are obligate intracellular parasites and death of the host cell halts viral replication. The context specific manner by which TNF acts indicates how the regulation of cellular decision strategies has evolved to provide specific physiological benefits.

Cellular signaling networks are often regarded as information processing units ^31–35^. However, the degree to which this is only metaphoric or is a faithful representation of biochemical processes is unclear. Our work shows that cell fate decisions are subject to the same fundamental principle, the speed-accuracy tradeoff, that governs many other complex decision making systems ^36^ including social insects ^37,38^, humans ^25,26^, and artificial neural networks ^39^. The applicability of such a universal principle to cellular signaling supports the view that these networks are indeed bona-fide information processing systems.

Tunability in apoptotic decision strategy can have two potential effects. First, it allows cells to respond dynamically to pathogen threat. In the work presented here, we did not vary the inflammatory signal over time. However, in the context of a true infection, it is likely that macrophages dynamically adapt to changes in pathogen load. Tunability in the decision strategy will allow cells to dynamically respond to such changes, adding another layer of feedback control. Second, different tissues likely have variable degrees of pathogen exposure and intrinsic tolerance to damage. The tunability of cellular decision strategies allows adapting cellular decision to specific tissue context. Extending the connection between immune regulation of individual cell decision making strategies and organ level phenotypes to other tissues would pave the way to a more unified theory of tissue immunology.

## Supplementary Movie

Movies are available on Figshare:

http://doi.org/10.6084/m9.figshare.12003534

**Supplementary Movie 1: HSV-1 spread with Activated macrophages.** 3T3 fibroblasts were co-cultured with 50% activated, inflammatory macrophages and infected with MOI 1 of HSV-1. Cells were imaged continuously for around 3 days. Left panel: Viral spread, HSV-1 shown in purple and cell nuclei in cyan. Right panel: Viral spread from left panel color-coded based on time of infection. Scale bar, 100μm.

**Supplementary Movie 2: HSV-1 spread with Naive macrophages.** 3T3 fibroblasts were co-cultured with 50% naive macrophages and infected with MOI 1 of HSV-1. Cells were imaged continuously for around 3 days. Left panel: Viral spread, HSV-1 shown in purple and cell nuclei in cyan. Right panel: Viral spread from left panel color-coded based on time of infection. Scale bar, 100μm.

**Supplementary Movie 3: HSV-1 spread with Activated TNF KO macrophages.** 3T3 fibroblasts were co-cultured with 50% activated, inflammatory TNF KO macrophages and infected with MOI 1 of HSV-1. Cells were imaged continuously for around 3 days. Left panel: Viral spread, HSV-1 shown in purple and cell nuclei in cyan. Right panel: Viral spread from left panel color-coded based on time of infection. Scale bar, 100μm.

**Supplementary Movie 4: HSV-1 spread in the absence of macrophages.** 3T3 fibroblasts were infected with MOI 1 of HSV-1. Cells were imaged continuously for around 3 days. Left panel: Viral spread, HSV-1 shown in purple and cell nuclei in cyan. Right panel: Viral spread from left panel color-coded based on time of infection. Scale bar, 100μm.

**Supplementary Movie 5: Light-sheet imaging of HSV-1 spread in whole, live mouse corneas.** R26-H2B-mCherry mouse corneas were infected with HSV-1 embedded in agarose and imaged for 48 hours. Scale bar, 100μm.

**Supplementary Movie 6: Light-sheet imaging of HSV-1 spread in whole, live TNF-treated mouse corneas.** R26-H2B-mCherry mouse corneas were infected with HSV-1, embedded in agarose with 50ng/ml TNFα and imaged for 48 hours. Scale bar, 100μm.

## Methods

### Cell Culture

Mouse 3T3 fibroblasts were cultured in DMEM supplemented with 10% newborn calf serum (NBCS), 2mM L-glutamine, 10U/ml Penicillin, and 10μg/ml Streptomycin and incubated at 37°C with 5% CO_2_.

### Mice

TNF^+/+^-RelA-Venus and TNF^−/−^-RelA-Venus mice (C57Bl/6 background) were provided by Dr. Alexander Hoffmann (UCLA) and used as a source of BMDM. Rosa26-H2B-mCherry hemizygous mouse embryos were purchased from the Riken Laboratory of Animal Resource Development and Genetic Engineering, re-derived at the University of California, Irvine Transgenic Mouse Core Facility, and maintained in the UCLA animal facility. C57Bl/6 J mice were purchased from the Jackson Labs. Both male and female mice aged 8-15 weeks were used in experiments. All animal experiments were approved by the UCLA Animal Research Committee and mice were maintained at an AALAC accredited animal facility in SPF conditions.

### Generation and activation of mouse BMDM

BMDM were differentiated from bone marrow incubated in DMEM with 10% fetal calf serum (FCS), 2mM L-glutamine, 10U/ml Penicillin, 10μg/ml Streptomycin, and 10ng/ml M-CSF (GenScript) for 7 days. BMDM were activated by priming overnight with 10nM IFNɣ (Peprotech), followed by 5 hours with 200ng/ml LPS (Invivogen).

### Cell stimulation and viral infection

12.5×10^4^ 3T3 cells were seeded per well of black 96-well plates in imaging media (Fluorobrite with 10% NBCS, 2mM L-glutamine, 10U/ml Penicillin and 10μg/ml Streptomycin) supplemented, where indicated, with dose titrations of recombinant mouse TNFα (Cell Signaling Technologies), 30μM Z-VAD(OMe)-FMK (Cayman Chemical), 30μM Necrostatin-1 (Sigma Aldrich), 0.25μg/ml Actinomycin D (ThermoFisher), or the specified doses of LCL-161 (Cayman Chemical). For live imaging, cells were labeled with 20ng/ml Hoechst 33342 (Life Technologies) and cell death was tracked by incorporation of either 1:10,000 Sytox Green (ThermoFisher) or 1:5000 Cytotox Green or Red (Essen Biosciences). Where indicated, cells were infected at the relevant MOI with Herpes Simplex Virus-1 strain KOS bearing either mCherry, tdTomato, or mCerulean-tagged VP26 small capsid proteins (provided by Dr. Prashant Desai, Johns Hopkins University). DMSO was added as a vehicle control where necessary.

To determine the rate of viral spread from neighbors, one batch of fibroblasts were infected overnight with MOI 6 HSV-1. Infected cells were washed, labeled with 1μM Cell Trace Far Red (ThermoFisher scientific) and co-cultured at low density with uninfected cells keeping the total number of cells at 1.6×10^4^ per well.

For TNFα pre-treatment experiments, cells were cultured overnight with dose titrations of TNFα, then washed 3x with PBS, and simulated with dose titrations of the synthetic STING agonist DMXAA (Invivogen) for 1.5h, or infected with MOI 1, 2, or 10 of HSV-1.

In macrophage-fibroblast co-culture experiments, 5h activated or naive BMDM were isolated, washed, and labeled with 1μM Cell Tracker Deep Red, Green, or Cell Trace Far Red (ThermoFisher scientific) for 10m at 37°C. Labeled cells were then co-cultured in black 96-well plates at variable density with 3T3 cells, keeping the total number at 1.6×10^4^ per well. When blocking TNF with neutralizing antibodies, anti-TNFα (Biolegend, LEAF-purified clone MP6-XT22) was used at 5μg/ml.

### Antibody staining

Cells were fixed on ice for 10m with 1.6% paraformaldehyde. Fixed cells were washed twice with PBS and permeabilized with ice cold 90% methanol. Fixed and permeabilized cells were washed 3x with PBS and blocked for 1h at room temperature with blocking buffer (3% BSA, 0.3% Triton-X-100 in PBS). The following primary antibodies were used: anti-Cleaved Caspase 8 (Cell Signaling Technologies, clone D5B2), anti-Cleaved Caspase 3 (Cell Signaling Technologies, clone D175), phospho-STING (Cell Signaling Technologies, clone D1C4T), anti-TNFα (Biolegend, clone MP6-XT22), anti-Ki-67 (Biolegend, clone 16A8). Cells were stained with primary antibodies diluted in blocking buffer overnight at 4°C. Primary stained cells were washed 3x with blocking buffer then incubated with secondary antibodies for 1h at room temperature, light protected. The following secondary antibodies were used: goat anti-Rabbit IgG-Alexa 594 or Alexa 647 (Jackson ImmunoResearch, cat 111-586-003, 111-606-003), goat anti-Rat IgG-Alexa 488 (Jackson ImmunoResearch, cat 112-546-003). Secondary stained cells were washed 3x with blocking buffer, twice with PBS, and counterstained with DAPI. To visualize TNF production by BMDM, cells were cultured for 5.5h with 10μg/ml Brefeldin A before fixation and staining.

For DISC proximity ligation experiments, cells were cultured on collagen and fibronectin coated glass slides, then fixed, permeabilized, and stained as above with the following antibodies: mouse anti-Caspase 8 (Proteintech, 2B9H8) and rabbit anti-Traf2 (Abcam, EPR6048). To perform proximity ligation, we used the DuoLink In Situ Proximity Ligation Assay Kit (Sigma Aldrich).

To detect apoptosis by Annexin V and FAM-FLICA-FMK, we used Annexin V-Alexa 568 (ThermoFisher scientific) and the Vybrant FAM Caspase 8 Assay kit (ThermoFisher scientific) according to manufacturer instructors.

### Generation of 3T3 Caspase 8 FRET Reporter Cells

The Caspase 8 FRET reporter plasmid pECFP-IETD2x-Venus ^24^ was cloned into a piggyBac destination vector with a Puromycin resistance cassette (pPB-ECFP-IETD2x-Venus-Puro) using Gateway Cloning. 3T3 cells were stably transfected with a piggyBac H2B-iRFP(710) plasmid and pPB-ECFP-IETD2x-Venus-Puro using Lipofectamine 3000. After 48h, double positive transfectants were selected using 2μg/ml Puromycin and Blasticidin. The brightest double positive cells were sorted by FACS (UCLA Broad Stem Cell Flow Cytometry Core) and expanded for experiments. For imaging, cells were cultured in collagen and fibronectin-coated glass-bottom 96-well plates.

### Quantification of gene expression by qPCR

2.5×10^5^ 3T3 cells were seeded per well of 6-well plates and stimulated with 50ng/ml TNFα. RNA was extracted and purified at the specified timepoints using the RNeasy Mini Kit (Qiagen). Complementary DNA was made using qScript XLT cDNA SuperMix (QuantaBio) and qPCR was performed using PerfeCTa SYBR Green MasterMix (QuantaBio) on a BioRad CFX96. The qPCR primer pairs were as follows:

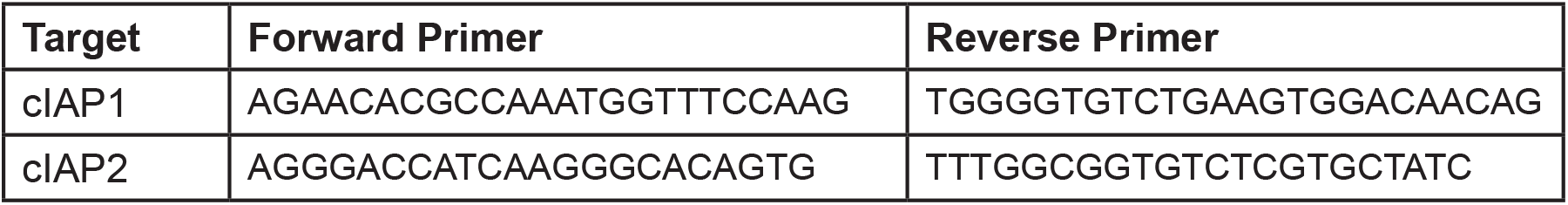

### Mathematical Modeling

The model is based on Gillespie’s algorithm, with interactions (infection) between neighboring cells. Cells in this model are scattered on a triangular grid. In our model, each cell can be in one of four states: (H) healthy, (I) infected, (D) dead following infection, or (B) dead but uninfected. The cells are organized on a 15×15 uniform triangular grid and the simulation is initialized by infecting the central cell on the grid. Each infected cell has a viral load that grows linearly and deterministically while the cell is alive and infected (Extended data 5C). Healthy cells can only be infected by live infected cells. The probability of infection scales with the total viral load of the cell’s nearest neighbors (Extended data 5D). The death rates of individual cells depend on their infection status and on their exposure to TNF, and are directly interpolated from measured values (Extended data 5E-F).

#### Model parameters

| Parameter | Meaning | Source |
| --- | --- | --- |
| VI | Viral infectivity | Extended data 5D |
| VGR | Viral growth rate | Extended data 5C |
| (TNF) | infected cell death rate | Extended data 5E |
| (TNF) | uninfected cell death rate | Extended data 5E |

#### Model Reactions

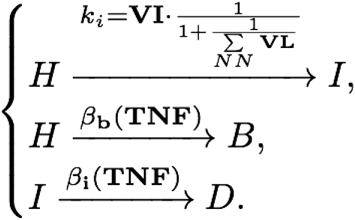

The viral load on an infected cell grows linearly

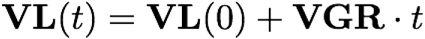

### Epifluorescence microscopy

Short-term or single time-point imaging was performed using either Nikon Plan Apo λ 10X/0.45 or 20X/0.75 objectives with a 0.7x demagnifier and Nikon Eclipse Ti microscope with a Flir, Chameleon3 CMOS camera. Long-term imaging was performed using a Carl Zeiss Plan-Apochromat 10x/0.45 objective with a 0.63x demagnifier and Carl Zeiss Axiovert 200m microscope also with a Flir, Chameleon3 camera housed inside an incubator for stable environmental control. All imaging was accomplished using custom automated software written using MATLAB and Micro-Manager. Image acquisition software is available on the GitHub repository: https://github.com/wollmanlab/Scope.

Automated image analysis was accomplished using custom-written software written in MATLAB. Image analysis software is available from the GitHub repository: https://github.com/wollmanlab/SingleCellVirusProcessing.

### Mouse cornea viral infection and embedding

Eyes were collected from R26-H2B-mCherry or C57Bl/6 mice and a small section of the corneal epithelium was disrupted using a 0.5mm rotating burr (Algerbrush, Gulden Ophthalmics). Whole eyes were then incubated overnight with 4×10^6^ PFU of HSV-1 mCerulean or tdTomato in pre-warmed cornea media (Fluorobrite with 10% FCS, L-glutamine, 10U/ml Penicillin, 10μg/ml Streptomycin, and 0.025μg/ml Amphotericin B). Whole eyes were washed 5x 5 minutes each in 10ml room temperature PBS. Corneas were then dissected and embedded in a 0.5ml syringe in a mixture of 1% low melting point agarose and cornea media supplemented with 1:50,000 Sytox Green and, where indicated, 50ng/ml TNFα. The embedded corneas were then briefly cooled at 4°C to solidify the agarose. The embedded cornea was then extruded from the syringe into an incubated chamber containing pre-warmed cornea media supplemented with Sytox Green and TNFα for imaging.

### Multi-color light sheet microscopy

We designed and built a high speed, multicolor light-sheet microscope specialized for large, live samples imaged over several days (Extended data 6A). Due to the large dimensions of our sample, an L-SPIM optical configuration was selected. Using the OpenSPIM design as a template ^40–42^, we modified several hardware components, and completely rewrote the acquisition software and methodology, to satisfy our unique demands.

#### Multicolor imaging

For multi-color illumination, we use an Omicron LightHUB laser combiner with 4 lasers:

1. Omicron LuxX 120mW 405nm laser
2. Omicron LuxX 100mW 488nm laser
3. Coherent OBIS 80mW 561nm OPSL
4. Omicron LuxX 100mW 638nm laser

We adapted the infinity-space tube of the OpenSPIM to house an optical filter changer (Sutter Instruments). We further modified the sample positioning arm of the 4D USB Stage (Picard Industries) to physically accommodate the filter wheel.

#### Modified sample chamber and objective

To allow a larger field of view, combined with enhanced resolution, we opted for a 16X 0.8NA Nikon CFI LWD Plan Fluorite Objective. For this, we modified the sample chamber, the sample chamber holder, and the objective holder ring from the original OpenSPIM design. For illumination, we used the Olympus UMPLFLN10XW objective.

#### Beam pivoting

To create a uniform excitation profile and minimize streaking artifacts, a 3khz resonant scanner (SC10-HF-6dia-20-3000, EOPC) was positioned along the beam path, perpendicular to the optical table plane. The scanner was run continuously during acquisition with an amplitude of 3 degrees.

#### Environmental control

Environmental control is achieved by positioning the entire microscope inside a Thermo Fisher Scientific Forma Steri-Cycle CO_2_ Incubator kept at 37°C with 5% CO_2_.

#### Stride-and-Strobe acquisition

To facilitate high-speed volume acquisition, we implemented a new acquisition mode we call Stride-and-Strobe (Extended data 6B). During imaging of a single volume, the sample is continuously moved with a velocity of 250μm/s in perpendicular to the imaging plane. While moving the sample, the camera is continuously imaging at a frame rate of 84 fps (11.85ms exposure). During that time, the sample makes a stride of 3μm. At the beginning of every individual frame exposure, a trigger is sent from the camera to the illumination laser. The laser turns on for a pulse of 2ms. The distance the sample moves during this exposure 0.5μm. The axial resolution of our microscope is 1.5μm. Therefore, this imaging strategy allows us to sample the tissue every 3μm while ensuring that motion-blur would not affect our image quality. Using this strategy, we can image a volume of 782 × 937 × 1500μm^3^ in roughly 6 seconds. To allow this mode of imaging, the Z-axis actuator of the 4D-Stage (Picard Industries) which moves the sample in the direction perpendicular to the imaging plane was replaced by a Hi-Res (0.75um step) Z-axis actuator (Picard Industries). To image a whole cornea, we first position the cornea in the transverse orientation in relation to the detection objective. We image the half of the cornea closest to the detection objective by tiling 18-25 image stacks with a minimal overlap of 15% of the FOV. We then rotate the cornea 90° to the en face orientation and image the half closest to the excitation objective in a similar manner. When multicolor imaging is applied, each block is imaged sequentially in different colors before continuing to the next block.

#### Image acquisition software

All imaging was accomplished using custom automated software written using MATLAB and Micro-Manager and is available through GitHub repository: https://github.com/wollmanlab/Scope

#### Data processing

Initial data processing was done using custom software implemented in Bash, ImageJ and Java (Extended data 6C). These steps relied heavily on the BigStitcher ^43^ ImageJ plugin. During acquisition, completed image stacks are transferred to a dedicated data analysis server. Upon arrival raw image stacks are converted into a multi-resolution HDF5 dataset. At the end of acquisition, the tiles are stitched together and the different acquisition angles (views) and channels are aligned as previously described ^44^. Blob-like objects like cells and beads are detected during this initial processing. In the case of time-lapse datasets, drift correction was applied. To allow for fast and efficient data processing, we implemented an automated and parallelized pipeline that incorporates these steps. Software is available through GitHub repository: https://github.com/alonyan/bigstitchparallel

#### Data presentation

Complete LSM datasets are fused as described in ^44^ and downsampled for presentation. Downsampled image stacks are rendered and presented using ImageJ and the 3DScript plugin ^45^ using custom ImageJ Macro scripts. For presentation, intensity levels for the bottom 5% are set to 0 (background). To account for bright outliers, intensity is scaled linearly so the top 0.01% (Nuclear channel) and 0.1% (other channels) of the pixels are saturated.

#### Data analysis

Data analysis is achieved using custom software implemented in MATLAB. Completed HDF5 datasets are accessed using the MIB software package.

## Acknowledgements

We thank Dr. Alexander Hoffmann (UCLA) for providing TNF KO mice, Dr. Prashant Desai (Johns Hopkins University) for providing fluorescent viruses, the CLICC Lux Lab (UCLA) for 3D printing of microscope components, and Zachary Hemminger for technical assistance. The work was funded by NIH grant R01EY024960 to RW.

---

**Extended Data 1.**
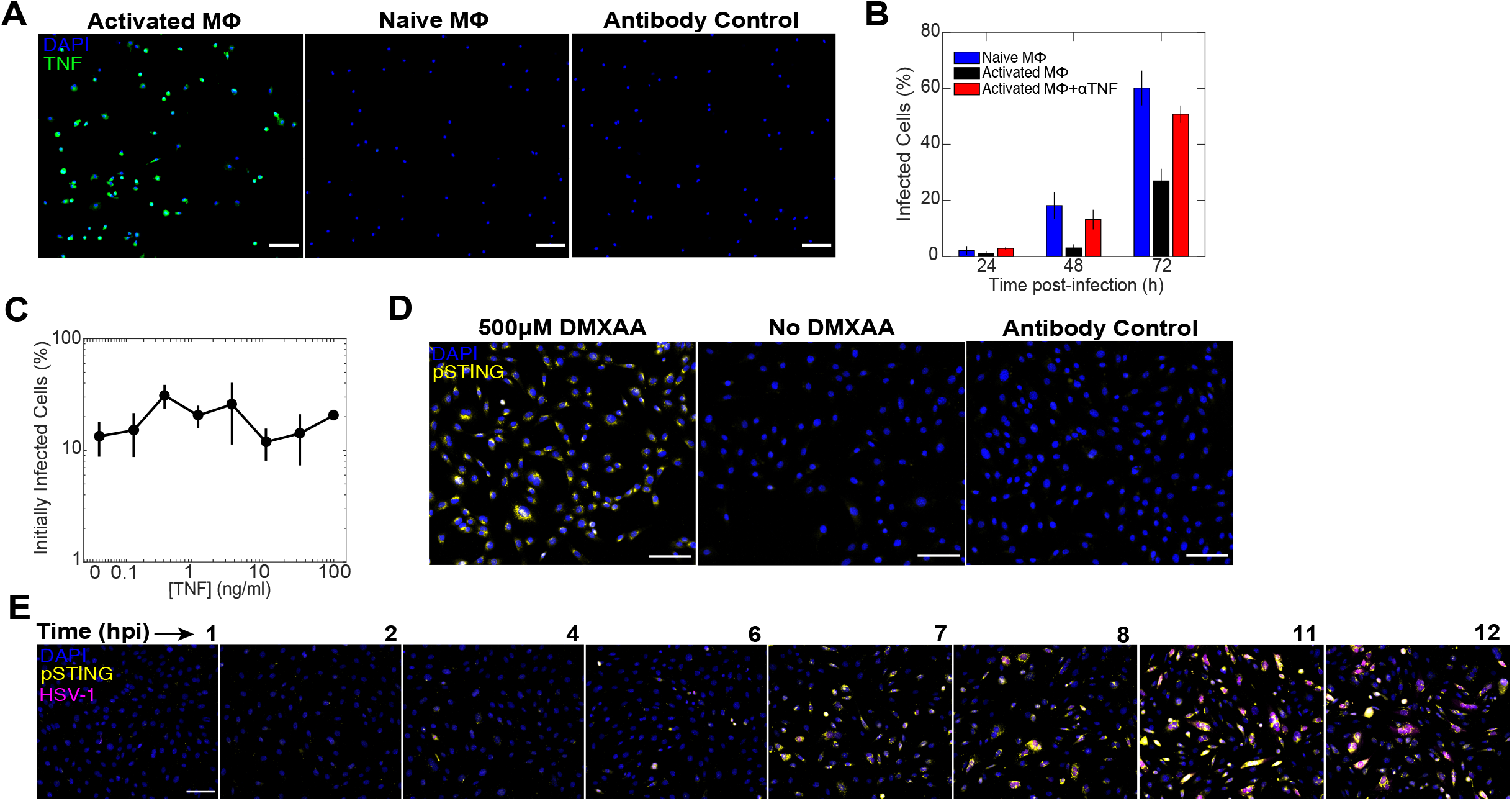
TNF production by activated macrophages restricts viral spread, but not through mechanisms of infectivity or viral sensing. **A**, Representative images of mouse BMDM activated with IFNγ and LPS, treated with Brefeldin A, and stained with anti-TNF. **B**, Naive or activated BMDM were co-cultured with fibroblasts, infected with MOI 1 HSV-1, and, where indicated, supplemented with neutralizing anti-TNF antibodies. The percentage of infected cells were quantified at 24, 48, and 72hpi. **C**, Quantification of cells infected only from virus in the media after being pulsed overnight with dose titrations of TNF and infected with MOI 2 of HSV-1. **D**, Representative images of fibroblasts stained for pSTING after being pulsed overnight with TNF, washed, and exposed to the synthetic STING agonist DMXAA. **E**, Representative images of fibroblasts stained for pSTING after being pulsed over-

**Extended Data 2.**
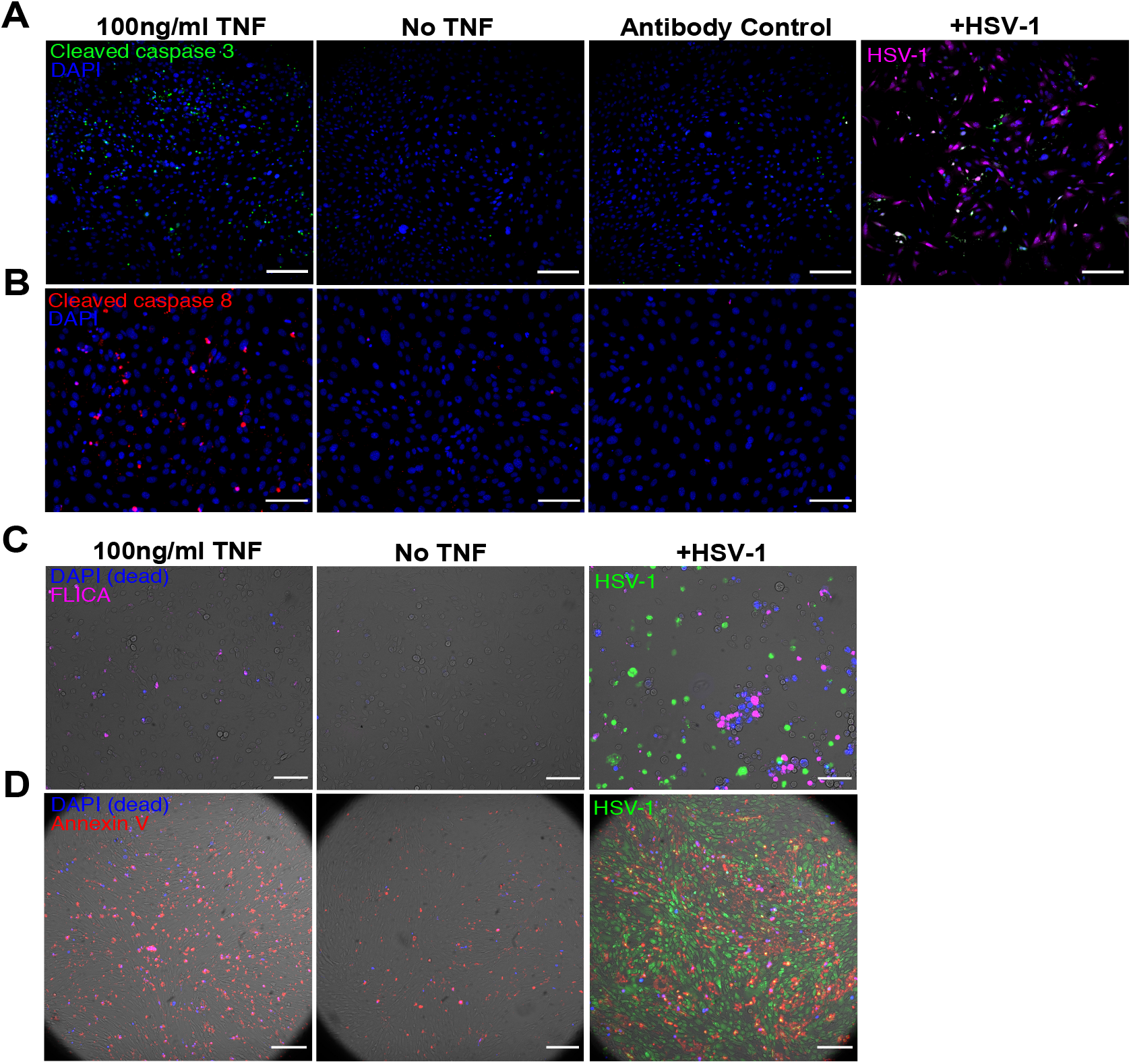
TNF-treated infected and uninfected cells die through apoptosis. **A**, Representative images of cells stained for cleaved Caspase 3 24h after TNF treatment or infection with MOI 6 of HSV-1. **B**, Representative images of cells stained for cleaved Caspase 8 24h after TNF treatment. **C**, Representative images of cells assayed for caspase activity by FLICA staining 24h after TNF treatment or infection with MOI 10 HSV-1. **D**, Representative images of cells stained for externalized phosphatidylserine by Annexin V staining 24h after TNF treatment or infection with MOI 6 of HSV-1. All scale bars,100μm.

**Extended Data 3.**
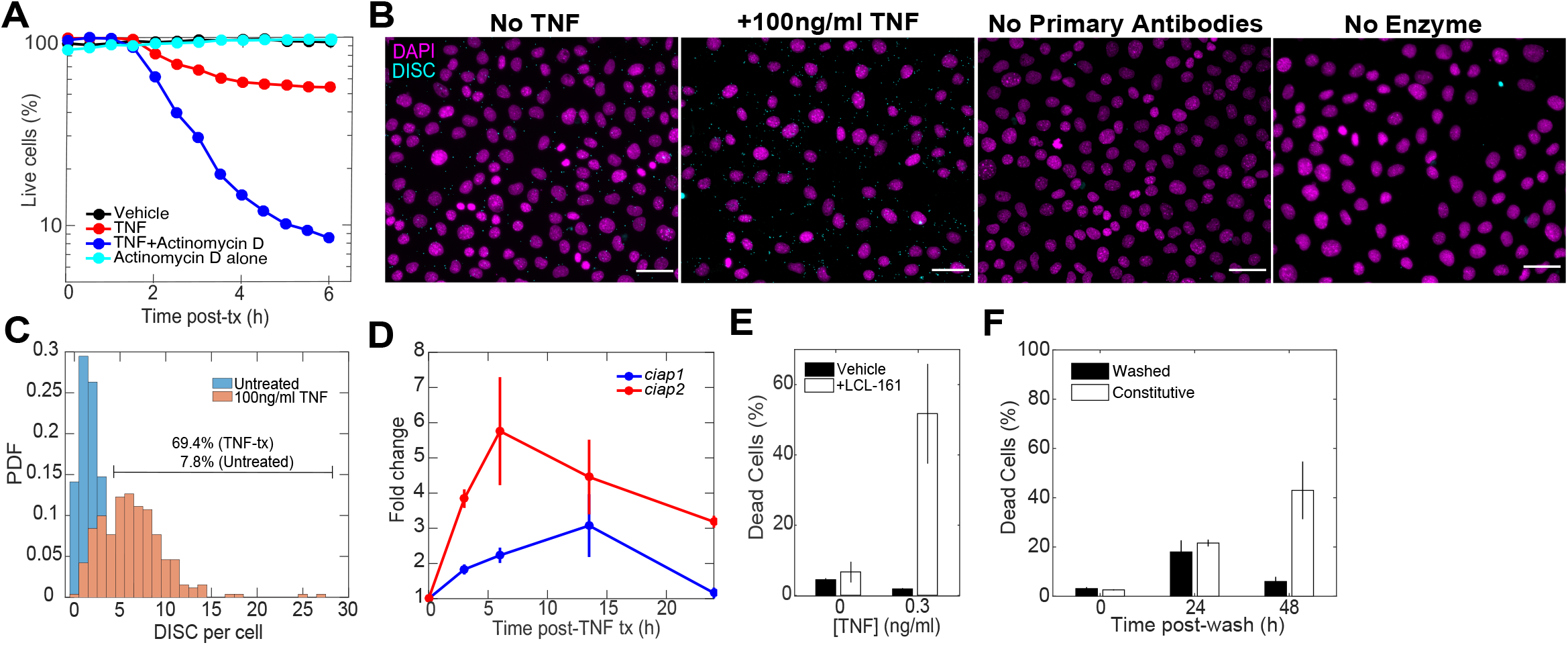
TNF transitions cells into a primed to death cell state. **A**, Quantification of cell death after treatment with TNF or TNF supplemented with Actinomycin D. **B**, Representative images of cells assayed for DISC assembly by Traf2-Caspase 8 promixity ligation. **C**,Quantification of number of DISC spots per cell for untreated and TNF-treated cells. Line shows the gate selected to distinguish positive cells. Number of cells = 475(Untreated), 261(TNF-tx) **D**, Quantification of TNF-induced *ciap1* and *2* by qPCR. **E**, Quantification of cell death for cells treated with 0 or 0.3ng/ml TNF and vehicle or 50pg/ml LCL-161. **F**, Quantification of cell death for cells treated constitutively or overnight with 10ng/ml TNF. All scale bars=100μm. **D-F** Data are mean +/− s.d. All scale bars,100μm.

**Extended Data 4.**
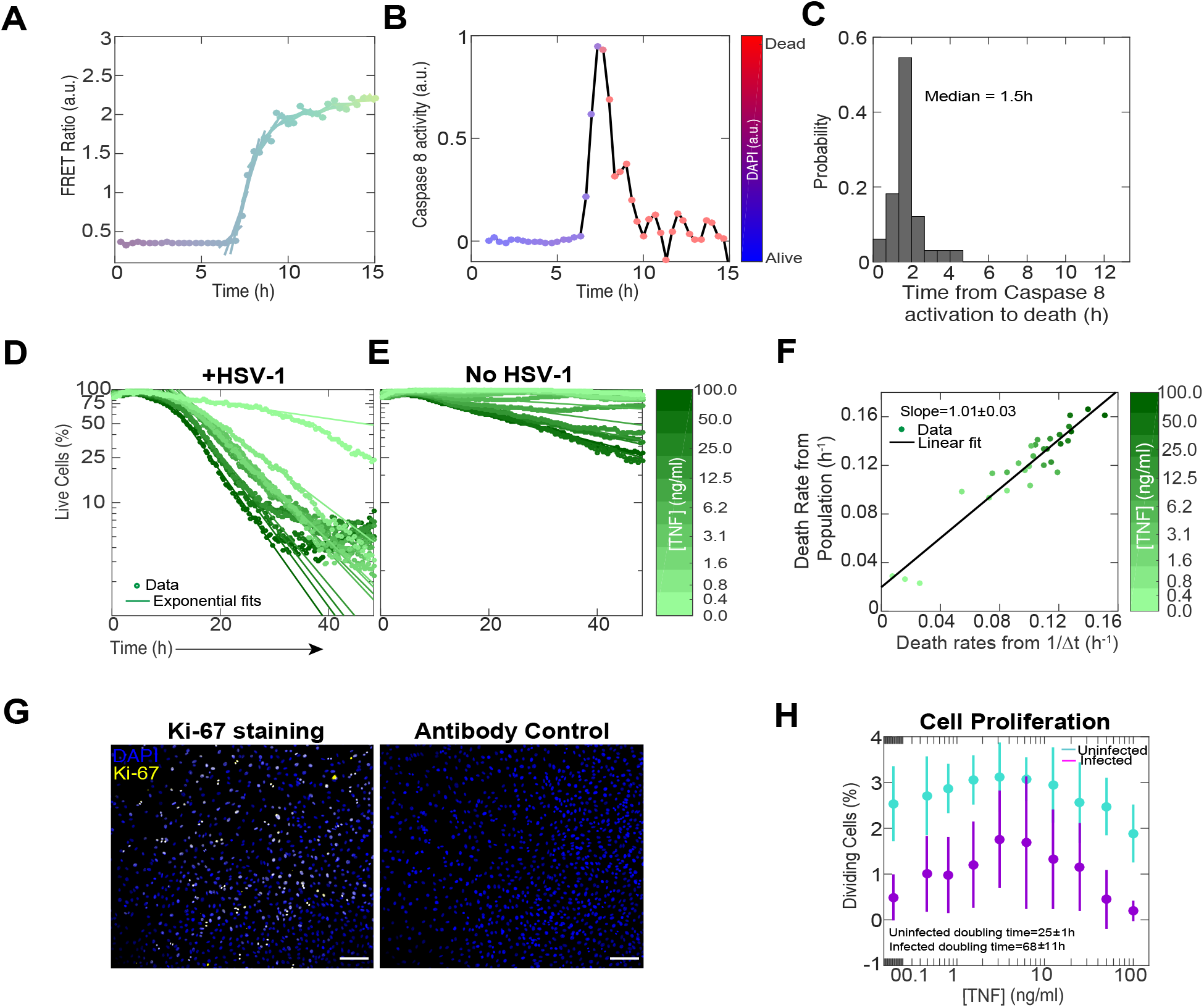
TNF causes the emergence of a speed versus accuracy tradeoff in the cellular decision to undergo apoptosis. **A,** Time courses of cleavage for Caspase 8 activity biosensor. Lines are piecewise linear-fits over a time window of ±2 frames around each timepoint. **B.** Caspase-8 activity measured as the slopes of the lines in panel **A.** Marker color indicates cell viability as measured using DAPI. **C,** Quantification of the time between the onset of Caspase 8 activity and death. Onset of Caspase 8 activity was determined by applying a 1d step filter and selecting the time of maximal response. **D**, Quantification of cell survival over time for cells treated with dose titrations of TNF and infected with MOI 10 HSV-1 or **E**, left uninfected. Points are data and solid lines represent exponential fits. **F**, Comparison of death rates quantified from exponential fits of population decay data versus single cell tracking. Points are data and solid line represents a linear fit. **G**, Representative images of cells stained for proliferation using Ki-67. **H**, Quantification of cell proliferation using Ki-67 staining for cells treated with TNF and infected with MOI 10 HSV-1. **H,** Data are mean +/− s.d. All scale bars,100μm.

**Extended Data 5.**
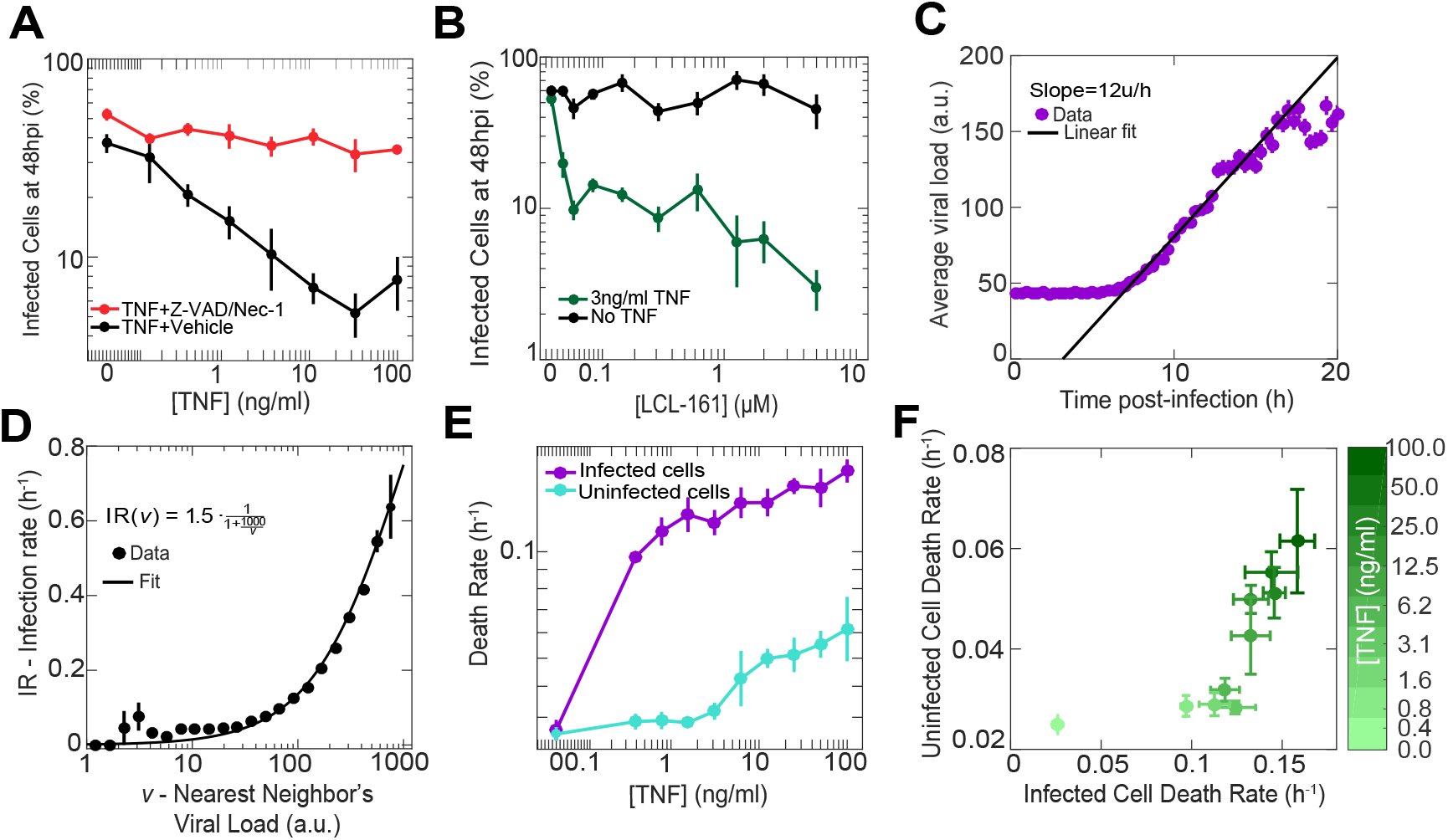
Experimental parameterization of mathematical model. **A**, Quantification of infected cells after treatment with dose titrations of TNF supplemented with vehicle or 30μM Z-Vad and 30μM Necrostatin-1. Cells were infected with MOI 1 of HSV-1. **B**, Quantification of infected cells after treatment with 0 or 3ng/ml TNF supplemented with dose titrations of LCL-161. Cells were infected with MOI 1 of HSV-1. **C**, Average viral load per cell quantified from HSV-1 fluorescence after infection with MOI 10 HSV-1. **D**, Quantification of the infection rate for cells based on the viral load of their nearest neighbor. **E**, Death rates for cells treated with dose titrations of TNF and infected with MOI 10 of HSV-1 or left uninfected. **F**, Quantification of the coupling between infected and uninfected cell death rates. **A-C, E-F,** Data are mean +/− s.d. **D**, Data are mean+/− s.e.m.

**Extended Data 6.**
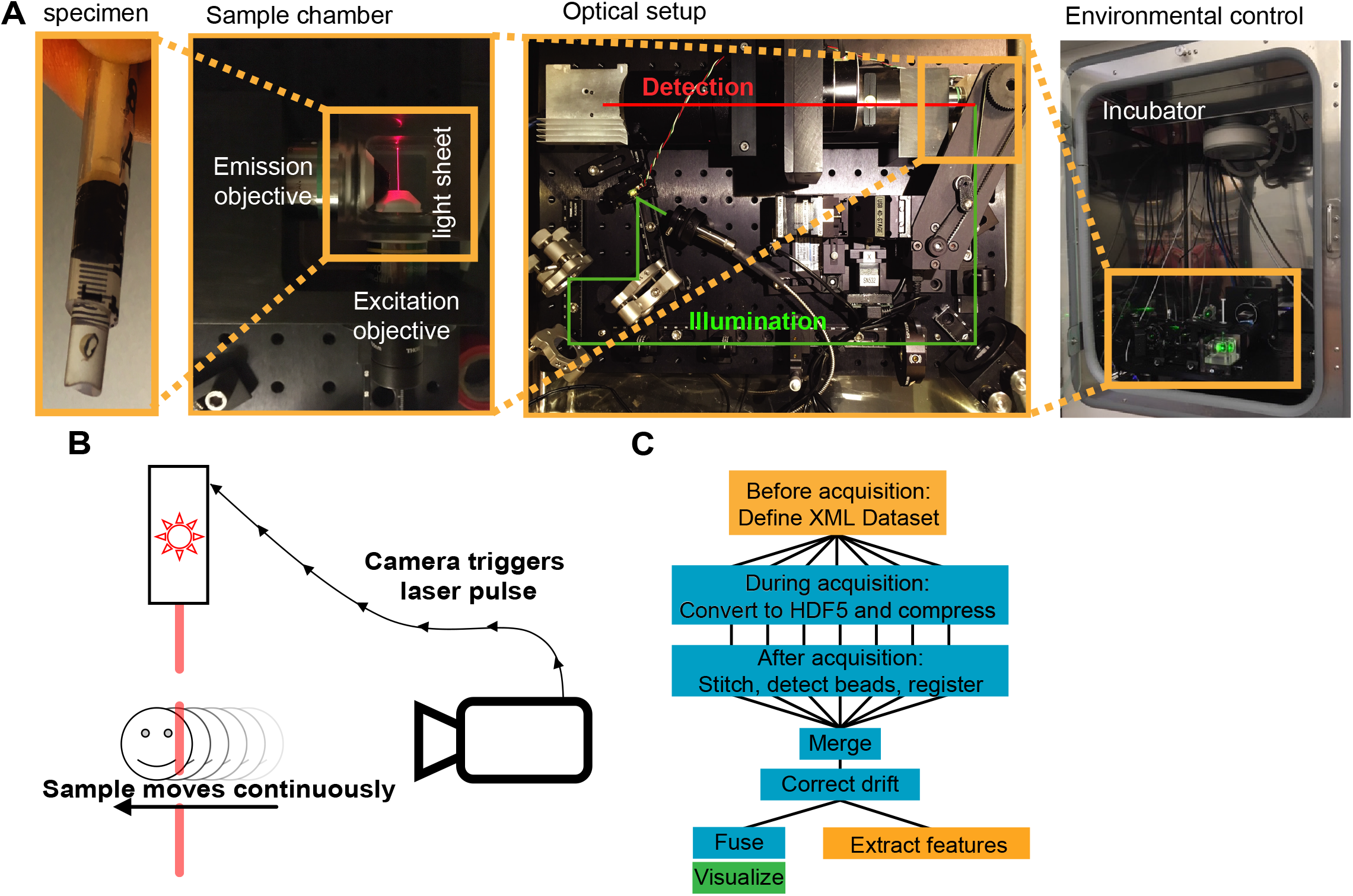
Custom light sheet microscopy setup. **A,** Specimens are embedded in a 0.5ml syringe in a mixture of 1% low melting point agarose and cornea media. The sample is mounted on a 4-D translation-rotation stage and submerged in a sample chamber for imaging. The original optical setup of the Open-SPIM was modified by adding a resonant mirror that provides beam pivoting for even illumination, an optical filter wheel to allow multichannel acquisition, and an alternative emission objective. The whole setup is mounted inside a tissue culture incubator for environmental control. **B,** To allow fast acquisition, the sample is translated continuously through the light sheet while the camera is acquiring images. At each frame, the camera triggers a short laser pulse. The mechanical stage was upgraded to allow finer translation along the axis of motion. **C,** Image acquisition and analysis pipeline. Orange boxes implemented in MATLAB, blue boxes use ImageJ and the BigStitcher plugin and implemented as automated bash scripts, and green box implemented using the 3DScript ImageJ plugin.

**Extended Data 7.**
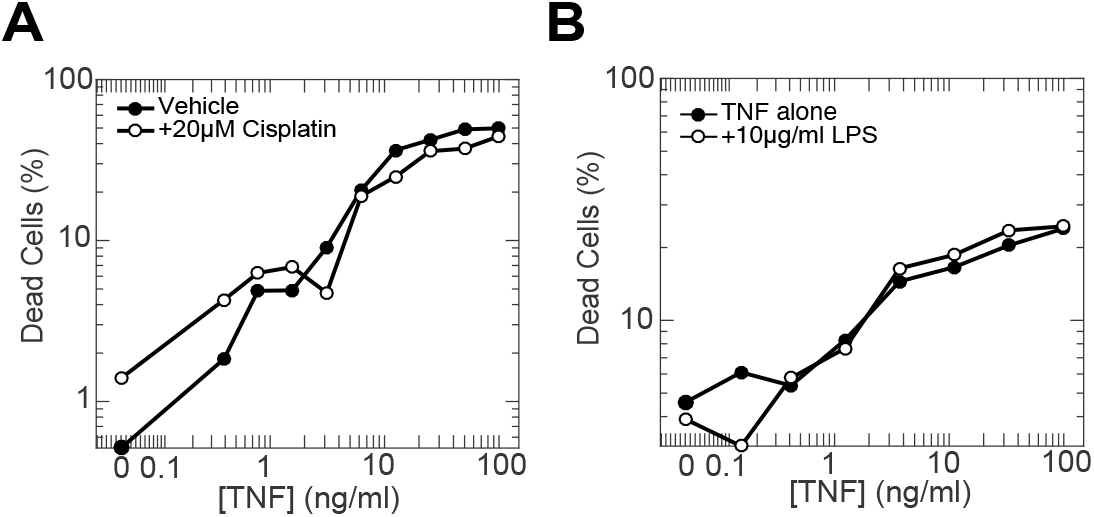
TNF-treatment does not sensitize cells to DNA damaging drugs or bacterial LPS. **A**, Cells were treated with dose titrations of TNF supplemented with either vehicle or 20μM Cisplatin. Cell death was quantified 48h post treatment. **B**, Cells were treated with dose titrations of TNF supplemented with either vehicle or 10μg/ml LPS. Cell death was quantified 24h post treatment.

## Notes

#### Summary of Updates

The text was revised for clarity and to include links to supp video. No change in the results or interpretation.

